# Astrocytic connection to engram neurons Increased after learning

**DOI:** 10.1101/2023.01.25.525617

**Authors:** Jooyoung Kim, Yongmin Sung, HyoJin Park, Dong Il Choi, Ji-il Kim, Hoonwon Lee, Min Kyo Jung, Seulgi Noh, Sanghyun Ye, Jiah Lee, Md Ariful Islam, Heejung Chun, Ji Young Mun, Bong-Kiun Kaang

## Abstract

Astrocytes directly participate in learning and memory. However, the structural association between astrocytes and memory-encoding engram neurons after learning remains to be elucidated. We developed astrocyte-enhanced green fluorescent protein reconstitution across synaptic partners (eGRASP) to examine tripartite synapses between astrocytes and engram neurons. Using astrocyte-eGRASP, we found that astrocytes had increased connections to engram neurons after learning. Dendritic spines with astrocytic contacts showed enhanced morphology. Live-cell imaging of astrocyte-eGRASP revealed that astrocytic connections are stabilized by neuronal activity. These results indicate that astrocytes distinguish contact between engram neurons and generate engram-specific contact patterns during learning.

## Introduction

Astrocytes are major participants in synaptic plasticity and memory. Astrocytes are glial cells that occupy a large percentage of central nervous system cells; however, for a long time, they have been regarded to play supportive functions in synaptic communication and cognition. Found in the last decade, astrocytes are involved in synaptic communication more directly (for review, Bazargani and Attwell, 2016). High-resolution imaging showed that a structure called tripartite synapses is the structural basis of such communication (for review, Araque et al., 1999; Perea et al., 2009). Cell type-specific manipulation tools have enabled further investigation of astrocyte-neuron communication, revealing that astrocyte activity is both necessary and sufficient for synaptic plasticity and memory encoding (Adamsky et al., 2018; Kol et al., 2020).

Memory is known to be encoded in an ensemble of neurons, called engram neurons (Liu et al., 2012), and its connections are called engram synapses (Choi et al., 2021; Choi et al., 2018). An engram, by definition, is the physical substrate of memory, which is activated during learning and reactivated during retrieval (for review, Josselyn et al., 2015; Josselyn and Tonegawa, 2020). Engrams can be identified by immediate early gene (IEG) expression, and IEG promoters are used to capture engram neurons in behavioral experiments (Roy et al., 2022). Engram neurons are necessary (Han et al., 2009) and sufficient (Ramirez et al., 2013) in various regions, including the hippocampus and amygdala for memory encoding.

Many memory studies have focused on the molecular, physiological, and morphological properties of engram neurons during the learning process (Choi et al., 2021; Choi et al., 2018; Han et al., 2007; Jeong et al., 2021; Lee et al., 2022). However, it has not been studied whether astrocytes, which have recently been defined as an important part of memory function, communicate with engram neurons owing to technical limitations. Because the small size of the astrocyte-neuron interface hinders observation under optical microscopy, the interaction between astrocytes and neurons has commonly been studied using high-resolution imaging technology, such as electron microscopy. Electron microscopy offers a nanostructure of the astrocyte-neuron interface (Risher et al., 2014); however, limitations exist in the examination of tripartite synapse dynamics in an engram-specific manner owing to limited labeling capacity and fixation process. In turn, fluorescent labeling of membranes has been used (Haber et al., 2006); however, it was relatively inaccurate because of the fluorescence intensity threshold and resolution limits.

Therefore, we developed a new tool for visualizing astrocyte-neuron contact, astrocyte-eGRASP, using eGRASP constructs (Choi et al., 2018), and verified it in vitro and in vivo. The eGRASP is a split fluorescent protein that reconstitutes across the contact site between the membranes. Previous studies (Choi et al., 2021; Choi et al., 2018) have utilized dual-color eGRASP to label two types of synapses. In this study, we modified eGRASP constructs to label the contact sites between astrocytes and neurons. Successful labeling of astrocyte-neuron contact dynamics was confirmed using live cell imaging of primary cultures and confocal and electron microscopy of brain slices. Using this tool, we investigated changes in the number of tripartite synapses in engram and non-engram neurons. We found that engram neurons made more contact with astrocytes after learning. Our study demonstrated that astrocytes communicate with engram neurons in distinct connection patterns.

## Materials & Methods

### Mice

All behavioral experiments were performed on 8-to 12-week-old C57BL/6N mice purchased from Samtako. Bio. Korea (Osan, South Korea). Mice were raised under a 12-h light-dark cycle in standard laboratory cages, and food and water were provided ad libitum. All procedures and animal care followed the regulations and guidelines of the Institutional Animal Care and Use Committee of Seoul National University. For primary culture, postnatal day 2 (P2) and embryonic day 17-18 (E17-18) C57BL/6N mice were purchased from Koatech (Pyeongtaek, South Korea).

### Construction of astrocyte-eGRASP

The glial fibrillary acidic protein (GFAP) promoter was inserted into dual-eGRASP DNA constructs as described in detail by Choi et al. (2018). For the labeling of astrocytes, mBeRFP was provided by Yang (Yang et al., 2013), and subcloned into GFAP-myriRFP-P2A-kSPOPTcSp32Nrx and GFAP-myriRFP-P2A-kSPOPTySp32Nrx.

### Adeno-associated virus production

Adeno-associated viruses serotype 1/2 (AAV1/2; AAV particle containing both serotype 1 and 2 capsids) was packaged as previously described (Choi et al., 2021; Choi et al., 2018). Briefly, AAV1/2 were purified from HEK293T cells that were transfected with plasmids containing each expression cassette flanked by AAV2 ITRs, p5E18, p5E18-RXC1, and pAd-ΔF6. AAV2/1 particles were collected using heparin-agarose suspensions (Sigma, cat. # H6508) or heparin-sepharose resin (Cytiva, cat. # 17099801). The titer was measured by quantitative RT-PCR.

### Primary culture

Primary astrocytes were prepared from P2 C57BL/6 mouse pups. The cerebral cortex was dissected from P2 mouse pups and was dissociated using 2.5% trypsin at 37 °C for 30 min and trituration. The dissociated single-cell suspension was plated in a six-well culture plate (SPL, cat# 30006). Cells were grown in an astrocyte culture medium (DMEM supplemented with 15% fetal bovine serum [FBS] and 1% penicillin/streptomycin [Hyclone, cat# SV30010]) at 37 °C in a 5% CO2 incubator. On the third day of culture, the cells were vigorously washed with pipetting, and the medium was replaced to remove other cell types.

For primary neuronal culture, hippocampal tissue was isolated from E17-18 C57BL/6 mouse embryos. The hippocampal tissue was dissociated using 2.5% trypsin at 37 °C for 30 min and trituration in plating media (MEM [Hyclone, cat# sh30024.01] with 10% FBS, 0.45% glucose, 1 mM sodium pyruvate, and 1% penicillin/streptomycin). Dissociated single-cell suspensions were plated in a 24-well culture plate (Thermo Scientific™, cat# 142475) coated with poly-D-lysine. Three hours after plating, the plating medium was removed and replaced with maintenance media (neurobasal medium [Gibco, cat# 21103049] with 2% B27 [Gibco, cat# 17504044], 2 mM GlutaMAX [Gibco, cat# 35050061], and 1% penicillin/streptomycin) at 37 °C in a 5% CO2 incubator.

After AAV transduction, primary astrocytes were detached by trypsinization and transferred to the primary neuronal culture.

### Live cell imaging

For live cell imaging, astrocyte-eGRASP-expressed primary cultures were imaged on an INCell Analyzer 2000(GE Healthcare, US) with 60x objectives. Cell culture was maintained at 37 °C and CO2 was supplied during the imaging. Images were obtained before and after bicuculline treatment. Images were obtained at 1-h intervals before bicuculline treatment to measure the basal state. After treatment with 10 μM bicuculline (Hellobio, cat# HB0893) or water for injection (WFI, Gibco, cat# A1287303) and incubation at 37°C in a 5% CO2 incubator for 40 min, images were recorded at the same location at 1-h intervals. We used the FIJI (Schindelin et al., 2012) HyperStackReg plugin (Sharma, 2018) to reconstruct and align images and used Imaris (Bitplane, Zurich, Switzerland) software to count astrocyte-eGRASP. The investigators who analyzed the images were unaware of the group and time points.

### Stereotaxic surgery

Mice (8–10 weeks old) were deeply anesthetized using ketamine/xylazine solution and positioned on a stereotaxic apparatus (Stoelting Co.). The viral mixture was injected into the target regions using a 31-G needle with a Hamilton syringe at a rate of 0.125 μL/min. The total injection volume per site was 0.5 μL, and the needle tip was positioned 0.05 mm below the target coordinate immediately before the injection for 2 min. After injection, the needle was left in place for an additional 7 min before being slowly withdrawn. The stereotaxic coordinates for dorsal CA3 were (AP: −1.7/ ML: ± 2.35/ DV: −2.4) and the coordinate for dorsal CA1 was (AP: −1.8/ ML: ± 1.45/ DV: −1.65 below the skull surface).

### Contextual fear conditioning

All mice were fear-conditioned 2–4 weeks after the AAV injection. Each mouse was caged 7 days before conditioning and habituated to the hands of the investigator and anesthesia chamber without isoflurane for 5 consecutive days. On the conditioning day, 250 μL of 5 mg/mL doxycycline solution dissolved in saline was injected intraperitoneally for 2-h before conditioning, under brief anesthesia using isoflurane.

For contextual fear conditioning, mice were placed in the fear conditioning chamber (Med Associates Inc.), and after 150 s, three electrical foot shocks were given (2 s, 0.75 mA). Approximately 30 s after the last foot shock, mice were returned to their home cages. One day after conditioning, the mice were placed in the conditioned context to measure their freezing behavior. Freezing behavior was recorded and scored using VideoFreeze software (Med Associates Inc.).

### Image analysis

We used the Imaris software (Bitplane, Zurich, Switzerland) to process the confocal images. Each image was cropped at the boundary of a single astrocyte by using the Imaris surface function. Each trackable fluorescent protein-labeled dendrite was manually denoted as a filament, during which other fluorescent signals were hidden to exclude any bias. Each cyan or yellow eGRASP signal was labeled as a cyan or yellow sphere.

For eGRASP density analysis, the number of denoted cyan or yellow spheres was manually counted along each denoted filament. Normalization of the eGRASP density was performed within each astrocyte using the average eGRASP on myriRFP-labeled dendrites.

### Immunogold label electron microscopy (Immuno-EM)

For immuno-EM, mouse brain slices were fixed by 2.5% glutaraldehyde and 2% paraformaldehyde in sodium cacodylate buffer (pH 7.2) at 4 °C. Samples were fixed in 1% osmium tetroxide (OsO4) for 30 min at 4 °C. The fixed samples were then dehydrated, performed using an ethanol gradient (50%, 60%, 70%, 80%, 90% and 100%) for 20 min each, and the samples were subsequently transferred to EM812 medium (Electron Microscopy Science, Hatfield, PA, USA). After impregnation with pure resin, specimens were embedded into the same resin mixture. Samples were sectioned (60 nm) with an ultramicrotome (Leica Ultracut UCT; Leica Microsystems, Vienna, Austria) and then collected on nickel grids. Post-embedding immunogold labeling was performed with anti-GFP (Invitrogen, #A-11120) and 9–11 nm colloidal gold conjugated to goat anti mouse IgG secondary antibodies (Sigma-Aldrich, #G7652). Following immunogold labeling, sections were double stained with UranyLess (EMS, #22409) for 2 min and 3% lead citrate (EMS, #22410) for 1 min. Sections were then analyzed using a transmission electron microscope at 120 kV (Tecnai G2, ThermoFisher, USA).

### Correlative Light and Electron Microscopy (CLEM)

Mice were deeply anesthetized with sodium-pentobarbital and intracardially perfused with 0.15 M cacodylate buffer before 2% paraformaldehyde and 0.5% glutaraldehyde in 0.15 M cacodylate buffer (pH 7.4). The pre-fixed brain was removed and stored in the same fixative overnight at 4 °C. Brain slices (150 μm thick) were cut on a vibratome (Leica, Germany) in ice-cold 0.15 M cacodylate buffer, and small pieces of hippocampal tissues were imaged using laser scanning confocal microscopy (Nikon, Japan) in z-stack mode within the mouse CA1 region. Confocal imaging was immediately followed by fixation 2.5% glutaraldehyde and 2% paraformaldehyde in preparation for EM. Tissues were post-fixed with 2% osmium tetroxide (OsO4)/1.5% potassium ferrocyanide (EMS, Sigma, USA) for 1 hour on ice and washed several times. Tissues were placed in 1% thiocarbohydrazide (TCH) (Ted Pella, USA) solution for 20 minutes and then placed in 2% aqueous OsO4 for 40 minutes. Thereafter, tissues were incubated in 2% uranyl acetate (EMS, USA) at overnight and lead aspartate solution at for 30 minutes to enhance membrane contrast. Tissues were dehydrated using a graded series of ethanol (20%, 50%, 70%, 90%, and 100%) on ice, infiltrated with acetone, mixture of resin and acetone, and pure resin. The resin was prepared from the Epon 812 kit (EMS, USA) following the manufacturer’s instructions. Samples are placed in embedding tubes with fresh Epon mixture at 60°C for 2 days. Embedded samples were trimmed, and 100 nm serial sections were cut from the block using an automated tape-collecting ultramicrotome (ATUM) (RMC, USA) to collect 600 sections with a diamond knife (Diatome, Switzerland) on carbon-coated Kapton tape. At each same serial position of EM Volume 110 μm × 110 μm × 30 μm was imaged by the ZEISS Atlas 5 software for Gemini 300 SEM (ZEISS, Germany) at the accelerating voltage of 5 kV. Image resolution in the xy plane was 5 nm/pixel. A total of 300 SEM images were fine-tune aligned using FIJI software (Schindelin et al., 2012) TrakEM2 plug-in (Cardona et al., 2012). As serial images, images that engulfment synapses of astrocytes were manually tracked and 3D-reconstructed by Reconstruct software (Fiala, 2005).

### Statistical analysis

All statistical analyses were performed using Prism 9 (GraphPad). Non-normal datasets were compared using a two-tailed Mann-Whitney test. The exact value of the sample size and statistical significance are reported in each figure legend.

## Results

To label astrocyte-neuron contacts, we modified our recent technique, dual-eGRASP (Choi, 2018, Science). Dual-eGRASP requires the expression of pre-and post-eGRASP compartments in presynaptic and postsynaptic terminals. For the in vitro test, we expressed pre-eGRASP in neurons and post-eGRASP in astrocytes for the eGRASP to reconstitute at the interface of astrocytes and neurons (Figure 1A-C). Each construct was expressed in primary culture using an AAV, and AAV-transduced primary neurons and astrocytes were co-cultured (Figure 1B). The neuronal membrane was labeled using myristoylated (myr) TagRFP-T (TRT), the astrocytic membrane was labeled with blue-excited RFP (mBeRFP), and astrocyte-neuron contact was labeled with cyan eGRASP (Figure 1C). In confocal imaging, cyan eGRASP labeled the interface between the neurons and astrocytes (Figure 1D). We named this technique astrocyte-eGRASP.

**Figure 1.**
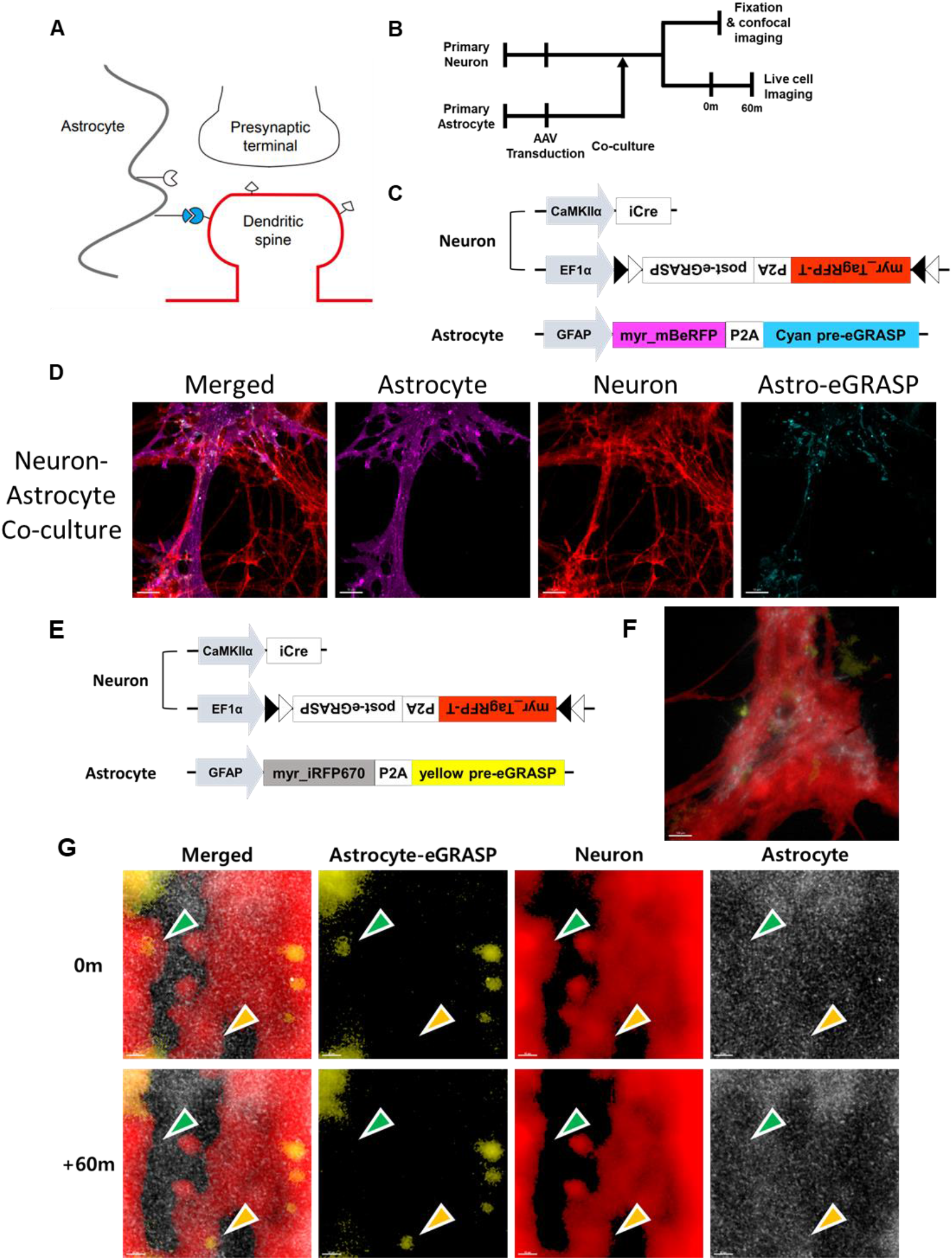
Visualization of neuron-astrocyte contact using eGRASP construct in vitro. (A) Schematic illustration for dendrite-astrocyte eGRASP. (B) Experimental procedure for in vitro astrocyte-eGRASP and live cell imaging. (C) Virus combination for in vitro astrocyte-eGRASP. (D) Representative image for in vitro astrocyte-eGRASP. Magenta: cyan-eGRASP expressing primary astrocyte. Red: post-eGRASP expressing primary neuron. Cyan: dendrite-astrocyte eGRASP. (E) Virus combination for live cell imaging of dendrite-astrocyte eGRASP. (F, G) Representative image for live cell imaging of dendrite-astrocyte eGRASP. Grey: yellow-eGRASP expressing primary astrocyte. Red: post-eGRASP expressing primary neuron. yellow: dendrite-astrocyte eGRASP. Green arrow: disappearing astrocyte-eGRASP puncta. Yellow arrow: appearing astrocyte-eGRASP puncta.

Astrocyte show fast morphological dynamics compared to neurons, and the interface of astrocytes and neuronal synapses is known to change within minute scale (Haber, 2006, J Neuro). To confirm whether astrocyte-eGRASP can represent astrocyte-neuron contact dynamics, we conducted live-cell imaging of the primary neuron-astrocyte coculture. The co-culture was prepared using the same method, except for the expression of myriRFP and yellow pre-eGRASP in astrocytes instead of mBeRFP and cyan pre-eGRASP (Figure 1E). Yellow astrocyte-eGRASP worked well, and a yellow fluorescent signal appeared at the interface between neurons and astrocytes (Figure 1F). The disappearance and appearance of astrocyte-eGRASP were observed at the surface of the neuronal membrane over a 1-h time interval (Figure 1G).

We then attempted to visualize tripartite synapses using astrocyte-eGRASP in vivo. Using dual-colored eGRASP, we labeled two of the three connections of the tripartite synapse (neuronal synapse, presynaptic terminal to astrocytic process, and postsynaptic terminal to astrocytic process) using two combinations of viruses (Figure 2). Each combination of AAV was injected into the hippocampal CA3 and CA1. In the first combination, we labeled the presynaptic terminal-to-astrocytic process with cyan-eGRASP and the postsynaptic terminal-to-astrocytic process with yellow eGRASP (Figure 2A). Thus, color-determining pre-eGRASP was expressed in the presynaptic and postsynaptic terminals, whereas post-eGRASP was expressed in astrocytes (Figure 2B). Consequently, yellow astrocyte-eGRASP appeared at the interface of myrmScarlet-I-labeled dendrites and myriRFP-labeled astrocytes, whereas cyan astrocyte-eGRASP labeled the interface between myriRFP-labeled astrocytes and axons (Figure 2C). In the second combination, we labeled neuronal synapses with cyan eGRASP and postsynaptic terminal-to-astrocytic process with yellow eGRASP. Color-determining cyan pre-eGRASP was expressed in CA3 presynaptic neurons, yellow pre-eGRASP was expressed in astrocytes, and post-eGRASP was expressed in CA1 postsynaptic neurons (Figure 2E). Cyan eGRASP appeared at the tip of the dendritic spine, whereas yellow eGRASP appeared on the surface of the dendrite at various locations (Figure 1F). To confirm whether astrocyte-eGRASP labels the astrocyte-neuron interface, ultrastructural analysis using electron microscopy was performed. Correlative light and electron microscopy (CLEM) revealed that astrocyte-eGRASP appeared at the interface of astrocytes and neuronal membranes (Figure 2G). In addition, immuno-gold labeling with anti-GFP confirmed the eGRASP located between astrocyte and neuron (Figure 2G).

**Figure 2.**
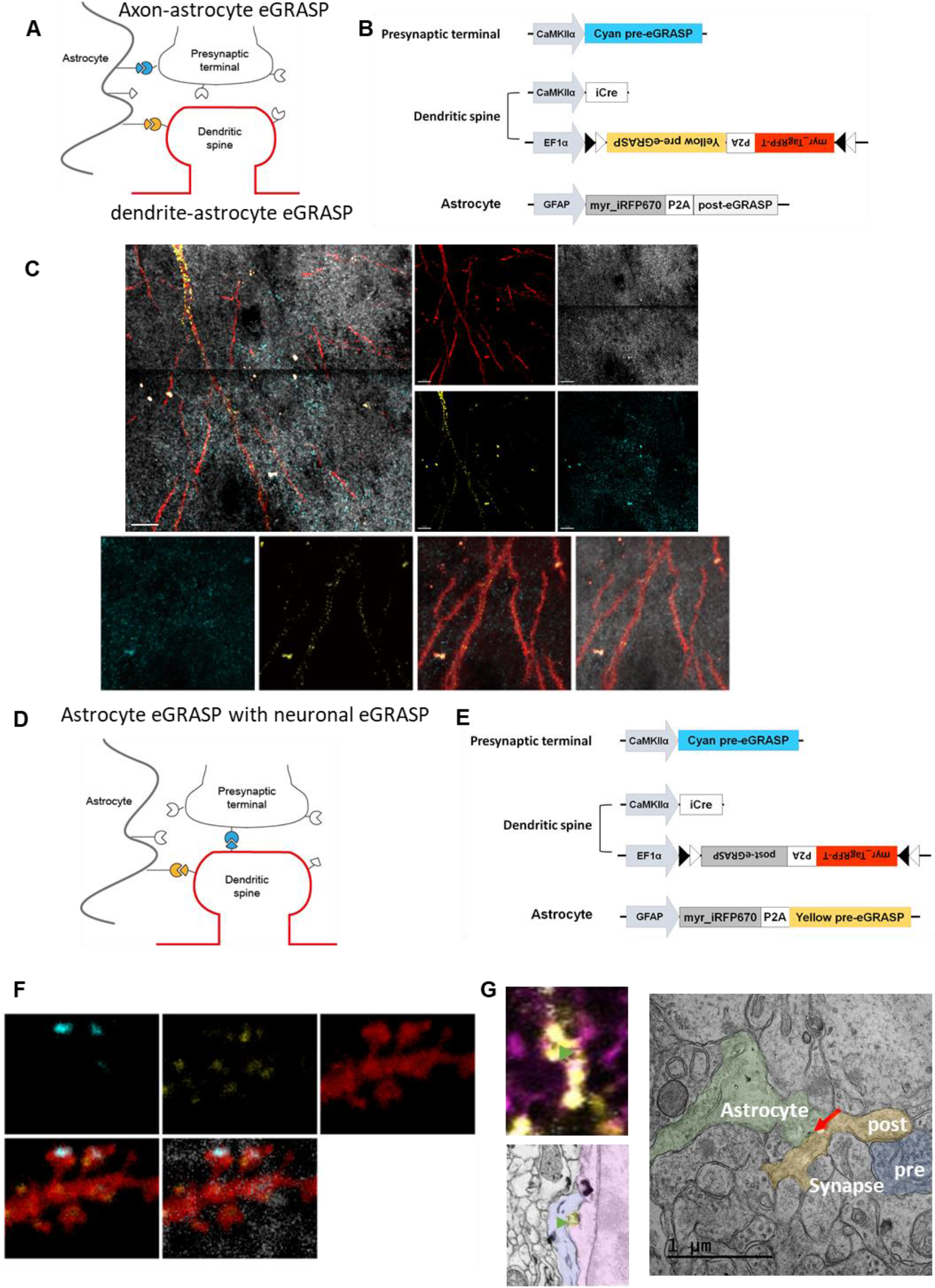
In vivo application of astrocyte-eGRASP. (A, B) Schematic illustration and viral combination for axon-astrocyte and dendrite-astrocyte eGRASP. (C) Representative image for axon-astrocyte and dendrite-astrocyte Grey: post-eGRASP expressing astrocyte. Red: yellow-eGRASP expressing CA1 dendrite. Cyan: axon-astrocyte eGRASP. Yellow: dendrite-astrocyte eGRASP. (D, E) Schematic illustration and virus combination for dendrite-astrocyte eGRASP with CA3-CA1 neuronal eGRASP. (F) Representative image for dendrite-astrocyte eGRASP with CA3-CA1 neuronal eGRASP. Grey: yellow-eGRASP expressing astrocyte. Red: post-eGRASP expressing dendrite. Cyan: CA3-CA1 neuronal eGRASP. Yellow: dendrite-astrocyte eGRASP (G) CLEM and immunogold EM images for astrocyte-eGRASP. Green arrow: astrocyte-eGRASP puncta in fluorescence and electron microscopic (EM) images. Red arrow: immunogold signal for astrocyte-eGRASP

To further evaluate astrocyte-neuron contact, we labeled the interface between CA1 dendrites and astrocyte processes with yellow eGRASP (Figure 3A, 3B) and performed a close examination of the astrocyte-eGRASP. While astrocyte-eGRASP appeared at the interface between myriRFP-labeled astrocytes and myrTRT-labeled neurons, it appeared not only at the tip of the dendritic spine but also at the shaft of the dendrite (Figure 3C). Synapses on dendritic shafts are mostly inhibitory synapses and, in a lesser proportion, excitatory shaft synapses (Muller et al., 2006; Reilly et al., 2011; Wable et al., 2014). A recent study using super-resolution microscopy reported astrocyte processes near GABAergic synapses in the hippocampal CA1 (Brunskine et al., 2022). We measured the ratio of astrocyte-eGRASP on spines and shafts, which showed that more astrocyte-eGRASP signals are on shafts at the dendritic trunk than at secondary or tertiary dendrites (Figure 3D). This is consistent with a previous report that more inhibitory synapses are on proximal dendrites than on distal dendrites in CA1 strata radiatum (Megias et al., 2001); although, we did not confirm whether the proportion of tripartite synapses differs according to dendrite type. Furthermore, we attempted to determine whether astrocyte-eGRASP can be applied to excitatory and inhibitory neurons of the CA1. We expressed yellow pre-eGRASP and myrTRT on excitatory neurons and cyan pre-eGRASP and myriRFP on inhibitory neurons using Cre recombinase and flippase, respectively (Figure 3E). Similar patterns of yellow eGRASP and cyan eGRASP puncta appeared at the surfaces of excitatory and inhibitory neuronal dendrites (Figure 3F). Astrocytes are known to play a role in inhibitory synapse formation (Elmariah et al., 2005); however, structural observation of tripartite synapses on inhibitory synaptic connections has rarely been reported because of technical difficulties. Therefore, astrocyte-eGRASP could be a strong tool that can be used to visualize not only the interaction of astrocytes on excitatory synaptic transmission but also inhibitory synaptic transmission.

**Figure 3.**
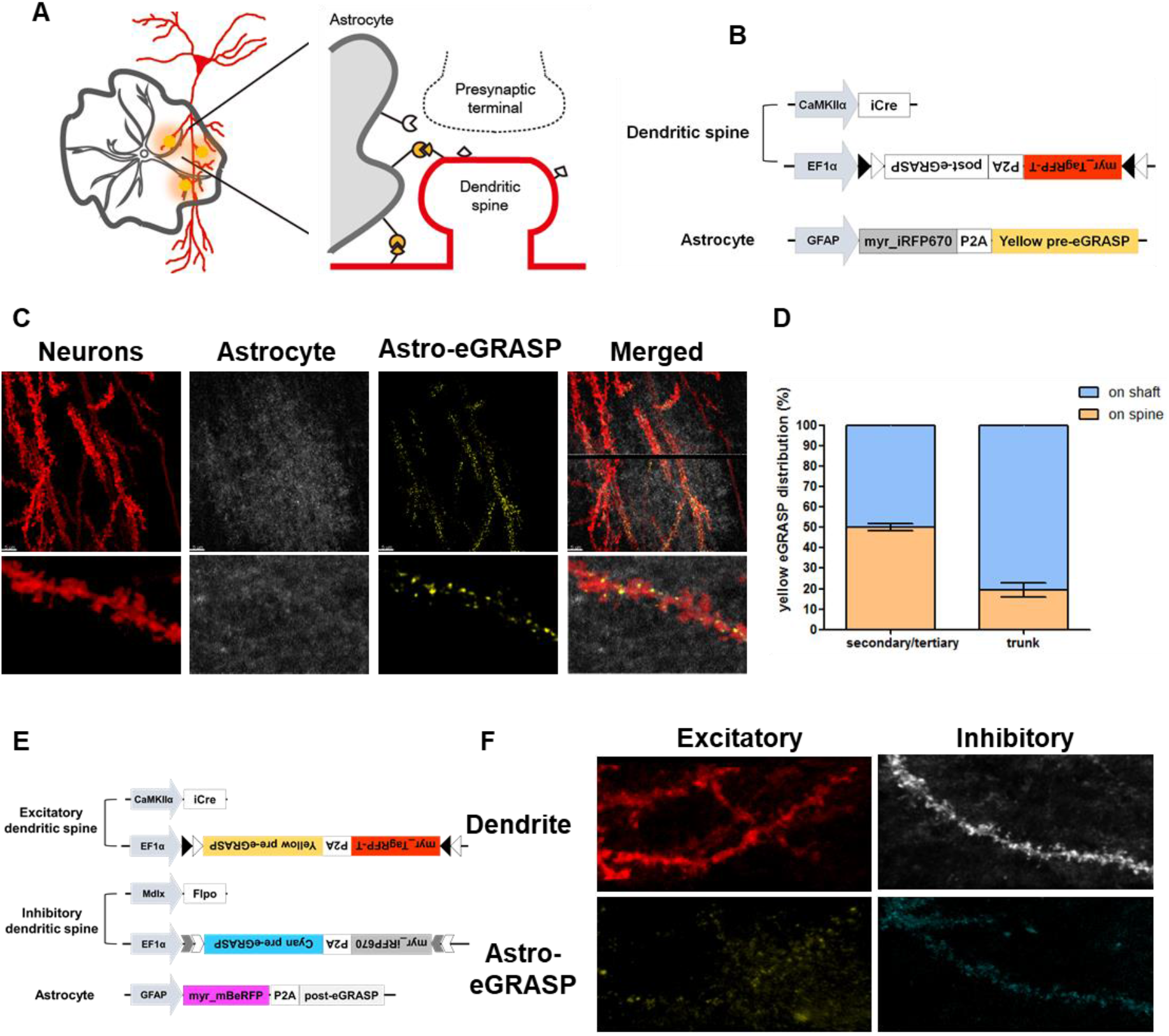
Astrocyte-neuron contacts on spines and shaft of excitatory and inhibitory neurons. (A, B) Schematic illustration and viral combination for astrocyte-eGRASP. (C) Representative image for astrocyte-eGRASP in CA1. Grey: yellow-eGRASP expressing astrocyte. Red: post-eGRASP expressing dendrite. Yellow: dendrite-astrocyte eGRASP. (D) Ratio of astrocyte-eGRASP on spine and shafts in secondary/tertiary dendrite or dendritic trunk. Secondary/tertiary dendrite, n =12; Trunk, n = 8. Data are represented as mean ± SEM. (E) Virus combination for astrocyte-eGRASP constructs expression at excitatory and inhibitory neurons. (F) Representative image for astrocyte-eGRASP on excitatory and inhibitory neurons. Red: yellow-eGRASP expressing dendrite of the excitatory neuron. Grey: cyan-eGRASP expressing dendrite of the inhibitory neuron. Yellow: dendrite-astrocyte eGRASP between excitatory neuron and astrocyte. Cyan: dendrite-astrocyte eGRASP between inhibitory neuron and astrocyte.

To determine the role of astrocyte connection to neurons in learning and memory, we applied astrocyte-eGRASP to the engram neurons of CA1 (Figure 4A, 4B). Engram neurons of CA1 were labeled with the IEG cfos-based tetO system. Doxycycline injection 2-h before contextual fear conditioning induced myrmScarlet-I and post-eGRASP expression in engram neurons. As an internal control, a random subset of excitatory neurons was labeled with myriRFP and post-eGRASP using Cre recombinase. Yellow pre-eGRASP was expressed in astrocytes and cyan pre-eGRASP was expressed in contralateral CA3 presynaptic neurons, as shown in Figure 2D. As the excitation/emission wavelengths for confocal imaging were mostly occupied, astrocytes were labeled with mBeRFP. Yellow astrocyte-eGRASP and cyan neuronal eGRASP appeared on the surface of engram and non-engram neurons, where they meet astrocytes (Figure 4C). Astrocyte-eGRASP appeared significantly more on the engram dendrites than on the non-engram dendrites, while the density of neuronal synapses from the contralateral CA3 did not differ between engram and non-engram neurons (Figure 4D). In addition, we measured the morphology of dendritic spines and analyzed them in the presence of astrocyte-eGRASP (Figure 4E). The dendritic spine with astrocyte-eGRASP showed a larger morphology than spines without astrocyte-eGRASP, both in engram and non-engram dendrites (Figure 4F). These results indicate that astrocytes specifically increase contact with engram neurons when memory is encoded, which are placed on more structurally potentiated dendritic spines.

**Figure 4.**
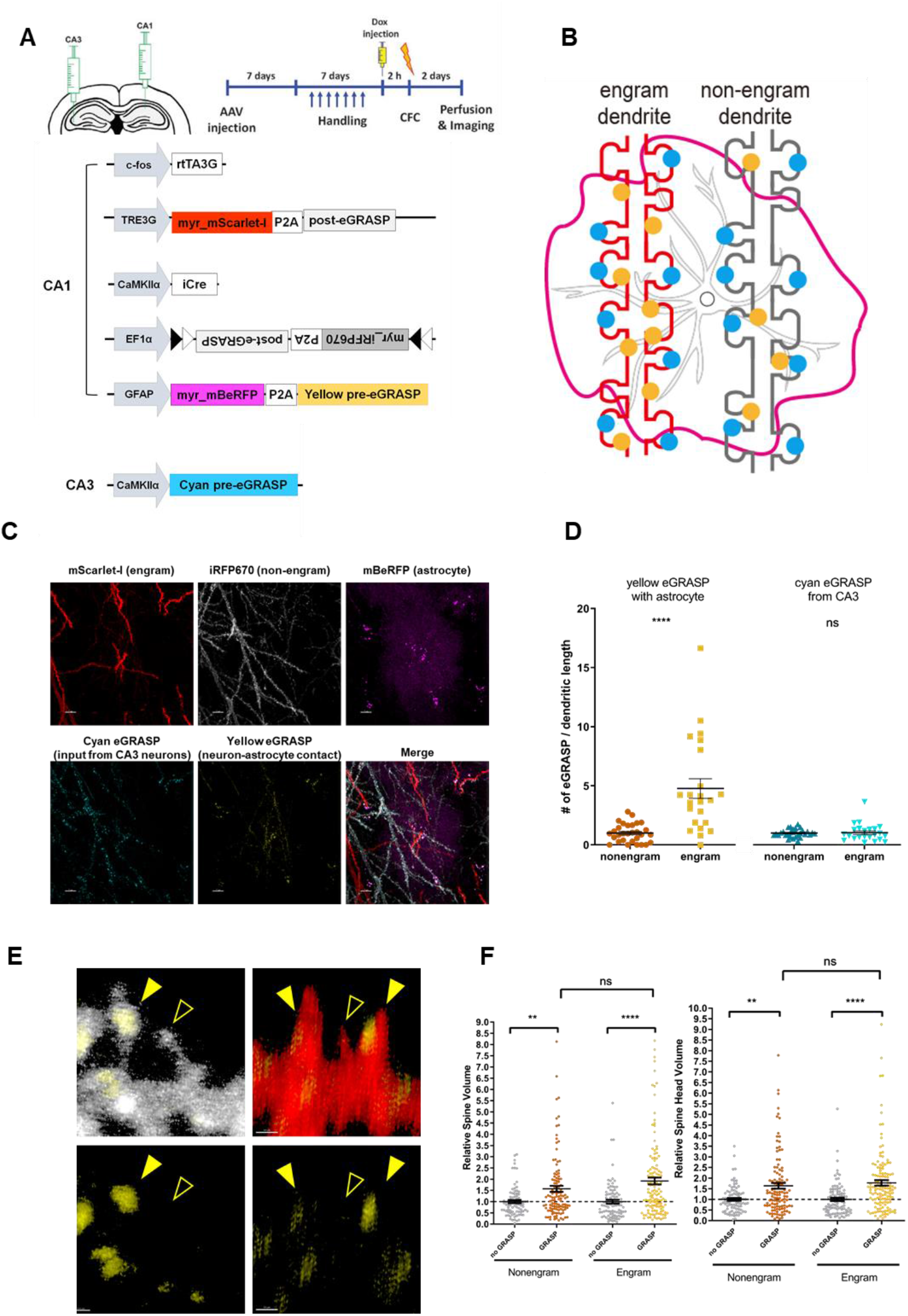
Increased astrocyte contacts on engram neurons. (A) Strategy and viral combination. (B) Schematic illustration for engram dendrite and non-engram dendrite with astrocyte-eGRASP (yellow) and CA3-CA1 neuronal eGRASP (cyan). (C) Representative images for engram neuron-astrocyte eGRASP. (D) Engram neurons have more astrocyte contact than non-engram neurons. Non-engram dendrite, n = 30; engram dendrite, n = 23. Mann-Whitney two-tailed test. n.s., not significant, ****p < 0.0001. Data are represented as mean ± SEM. (E) Representative images for dendritic spines with (filled arrow) or without (empty arrow) astrocyte-eGRASP in non-engram and engram neurons. (F) Dendritic spines with astrocyte-eGRASP have larger morphology. Non-engram, no eGRASP spine, n = 84; non-engram, eGRASP spine, n = 114; engram, no eGRASP spine, n = 93; engram, eGRASP spine, n =131. Mann-Whitney two-tailed test. n.s., not significant, **p < 0.01, ****p < 0.0001. Data are represented as mean ± SEM.

We then investigated the mechanism of astrocyte recognition and reaction to engram neurons. We conducted live cell imaging and analyzed time-lapse images of astrocyte-eGRASP in the primary co-culture during neuronal activation using bicuculline (Figure 5). Astrocyte motility is known to occur over minutes (Haber et al., 2006). Astrocyte-eGRASP dynamics were imaged with a 1-h time gap before and after bicuculline application (Figure 5A). Comparing the bicuculline-applied dynamics to the baseline, we found that increased neuronal activity reduced the disappearance of the astrocyte-eGRASP signal (Figure 5B), while the appearance of the astrocyte-eGRASP did not change (Figure 5C). In other words, the stability of astrocyte-neuron contacts increased after neuronal activity. This result suggests that the increase in astrocyte-eGRASP in engram neurons in vivo is mediated by the reduced disappearance of astrocyte-eGRASP after increased neuronal activity during learning.

**Figure 5.**
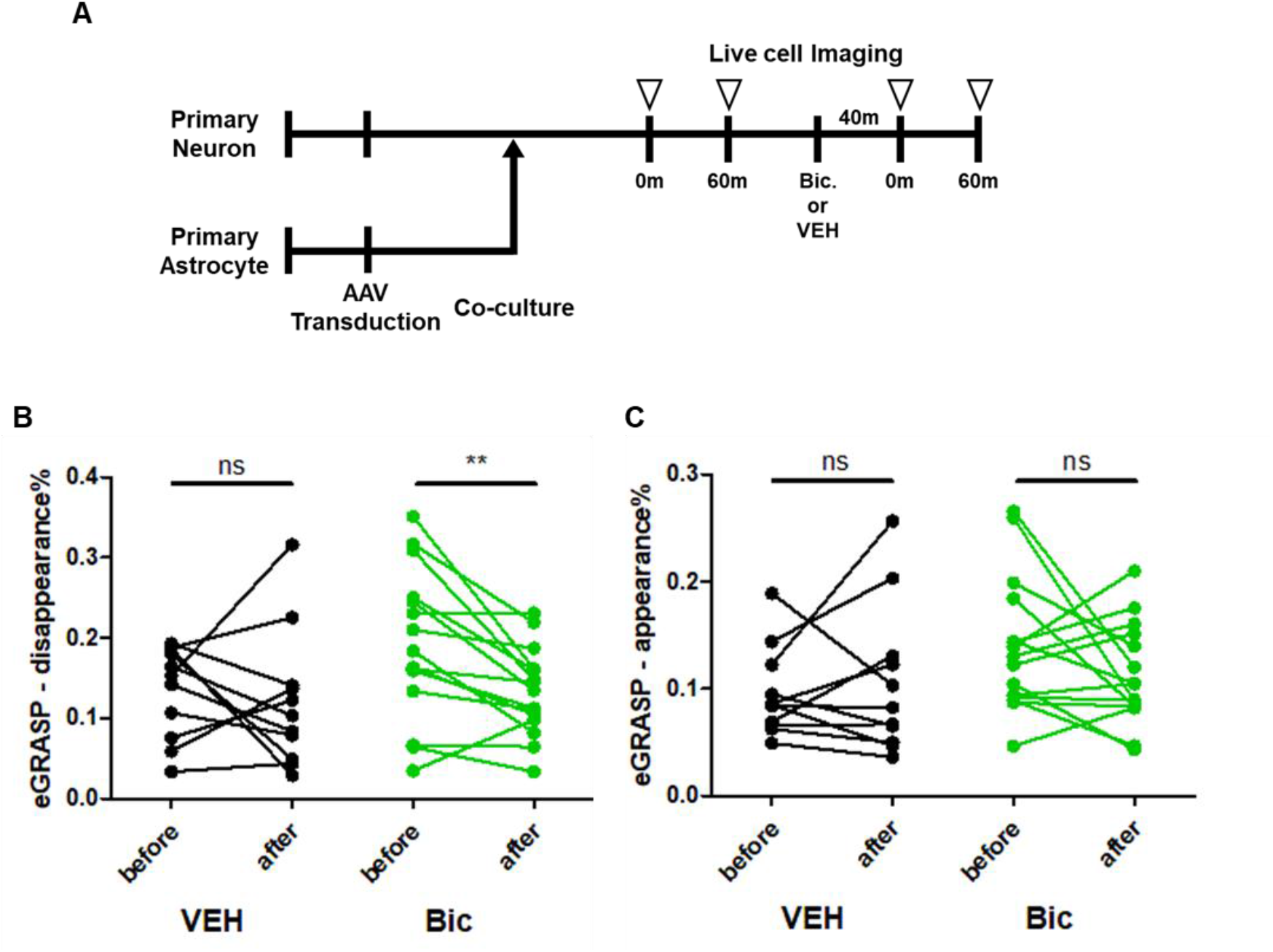
Increased stability of astrocyte-eGRASP after neuronal activation. (A) Schematics for the experiment. (B, C) Disappearance (B) and appearance (C) of astrocyte-eGRASP before and after the drug application. VEH for the vehicle-treated group and Bic for the bicuculline-treated group. Vehicle, n = 11; Bicuculline, n = 15. Two-tailed paired t-test. n.s., not significant; **p < 0.01.

## Discussion

Understanding the structure of astrocyte-neuron connections is key to understanding the function of astrocytes in synaptic communication and cognition. In the present study, we developed a new fluorescence labeling technique, astrocyte-eGRASP, that can visualize astrocyte-neuron connections. Astrocyte-eGRASP enabled us to monitor astrocyte-neuron connections both in vitro and in vivo and successfully labeled tripartite synapses when combined with neuronal eGRASP. Using astrocyte-eGRASP, we found that astrocytes establish more contact with memory-encoding engram neurons than with non-engram neurons. Furthermore, we demonstrated that neuronal activity increased the stability of astrocyte-neuron connections.

Here, we report that astrocyte connections to memory-encoding engram neurons are increased compared to non-engram neurons. During memory formation, synapses between engram neurons increase in number and size (Choi et al., 2018). Our results add to the finding that astrocyte participation is increased in engram neurons as well as in neuronal synapses. Furthermore, we found that spines with astrocyte contacts had a larger morphology. This is consistent with a previous report that astrocyte contact is more stable in spines with larger volumes (Haber et al., 2006), although whether astrocytes maintain spine maturation or whether spine maturation stabilizes astrocyte contact is unclear. It is conceivable that astrocytes make more contact with engram neurons to maintain the potentiated synapses of engram neurons that encode memory.

We found that single astrocyte can distinguish between engram and non-engram neurons and have distinct connectivity to engram neurons. This raises the question of how astrocytes sense the engram neurons. It is known that the engram neurons are activated during memory formation. Astrocytes have been reported to sense neuronal activity and respond to calcium signaling (Stobart et al., 2018). Here, we show that neuronal activity increases the stability of astrocyte contact. This result suggests that engram neuronal activity may affect astrocytes in the microenvironment and change astrocyte connectivity.

We showed that astrocytes make a considerable number of contacts with both dendritic spines and shafts. Tripartite synapses on dendritic spines are frequently reported and studied, but studies on tripartite synapses on the shaft are relatively rare (for review, Ishibashi et al., 2019). Shaft synapses consist of inhibitory synapses (Megias et al., 2001) and excitatory shaft synapses (Bucher et al., 2020). Electrophysiological studies have reported that astrocytes respond to inhibitory synaptic currents (Matos et al., 2018), but their role in engram formation and cognitive function is unclear. Excitatory synapses on the dendritic shaft are considered a sign of synaptogenesis (Bourne and Harris, 2011; Reilly et al., 2011), which is an important mechanism of the memory formation process (Lee et al., 2022). Astrocytes may participate in the shaft-to-spine transition of excitatory synapses (Risher et al., 2014), indicating that shaft astrocyte contact may play an important role in spine maturation. Future studies should investigate the role of shaft tripartite synapses in cognition and learning.

Here, we show that astrocytes make distinct and direct contact with memory-encoding engram neurons. Each astrocyte makes increased contact with engram neurons compared with non-engram neurons after learning. Moreover, astrocyte contact stabilizes after neuronal activity. Our results suggest that astrocyte participation in memory is mediated by enhancing tripartite synapses with engram neurons. These findings provide an important structural basis for studying astrocytic involvement in learning and memory. Future studies should determine the role of astrocytic contacts in engram neurons.

## Acknowledgement

This study was supported by the National Honor Scientist Program (NRF-2012R1A3A1050385) and KBRI basic research program by Ministry of Science and ICT (23-BR-01-03) of Korea.

## Notes

### Competing Interest Statement

The authors have declared no competing interest.

## References

Adamsky, A., Kol, A., Kreisel, T., Doron, A., Ozeri-Engelhard, N., Melcer, T., Refaeli, R., Horn, H., Regev, L., Groysman, M., et al. (2018). Astrocytic Activation Generates De Novo Neuronal Potentiation and Memory Enhancement. Cell 174, 59–71 e14.

Araque, A., Parpura, V., Sanzgiri, R.P., and Haydon, P.G. (1999). Tripartite synapses: glia, the unacknowledged partner. Trends Neurosci 22, 208–215.

Bazargani, N., and Attwell, D. (2016). Astrocyte calcium signaling: the third wave. Nat Neurosci 19, 182–189.

Bourne, J.N., and Harris, K.M. (2011). Coordination of size and number of excitatory and inhibitory synapses results in a balanced structural plasticity along mature hippocampal CA1 dendrites during LTP. Hippocampus 21, 354–373.

Brunskine, C., Passlick, S., and Henneberger, C. (2022). Structural Heterogeneity of the GABAergic Tripartite Synapse. Cells 11.

Bucher, M., Fanutza, T., and Mikhaylova, M. (2020). Cytoskeletal makeup of the synapse: Shaft versus spine. Cytoskeleton (Hoboken) 77, 55–64.

Cardona, A., Saalfeld, S., Schindelin, J., Arganda-Carreras, I., Preibisch, S., Longair, M., Tomancak, P., Hartenstein, V., and Douglas, R.J. (2012). TrakEM2 software for neural circuit reconstruction. PLoS One 7, e38011.

Choi, D.I., Kim, J., Lee, H., Kim, J.I., Sung, Y., Choi, J.E., Venkat, S.J., Park, P., Jung, H., and Kaang, B.K. (2021). Synaptic correlates of associative fear memory in the lateral amygdala. Neuron 109, 2717–2726 e2713.

Choi, J.H., Sim, S.E., Kim, J.I., Choi, D.I., Oh, J., Ye, S., Lee, J., Kim, T., Ko, H.G., Lim, C.S., et al. (2018). Interregional synaptic maps among engram cells underlie memory formation. Science 360, 430–435.

Elmariah, S.B., Oh, E.J., Hughes, E.G., and Balice-Gordon, R.J. (2005). Astrocytes regulate inhibitory synapse formation via Trk-mediated modulation of postsynaptic GABAA receptors. J Neurosci 25, 3638–3650.

Fiala, J.C. (2005). Reconstruct: a free editor for serial section microscopy. J Microsc 218, 52–61.

Haber, M., Zhou, L., and Murai, K.K. (2006). Cooperative astrocyte and dendritic spine dynamics at hippocampal excitatory synapses. J Neurosci 26, 8881–8891.

Han, J.H., Kushner, S.A., Yiu, A.P., Cole, C.J., Matynia, A., Brown, R.A., Neve, R.L., Guzowski, J.F., Silva, A.J., and Josselyn, S.A. (2007). Neuronal competition and selection during memory formation. Science 316, 457–460.

Han, J.H., Kushner, S.A., Yiu, A.P., Hsiang, H.L., Buch, T., Waisman, A., Bontempi, B., Neve, R.L., Frankland, P.W., and Josselyn, S.A. (2009). Selective erasure of a fear memory. Science 323, 1492–1496.

Ishibashi, M., Egawa, K., and Fukuda, A. (2019). Diverse Actions of Astrocytes in GABAergic Signaling. Int J Mol Sci 20.

Jeong, Y., Cho, H.Y., Kim, M., Oh, J.P., Kang, M.S., Yoo, M., Lee, H.S., and Han, J.H. (2021). Synaptic plasticity-dependent competition rule influences memory formation. Nat Commun 12, 3915.

Josselyn, S.A., Kohler, S., and Frankland, P.W. (2015). Finding the engram. Nat Rev Neurosci 16, 521–534.

Josselyn, S.A., and Tonegawa, S. (2020). Memory engrams: Recalling the past and imagining the future. Science 367.

Kol, A., Adamsky, A., Groysman, M., Kreisel, T., London, M., and Goshen, I. (2020). Astrocytes contribute to remote memory formation by modulating hippocampal-cortical communication during learning. Nat Neurosci 23, 1229–1239.

Lee, C., Lee, B.H., Jung, H., Lee, C., Sung, Y., Kim, H., Kim, J., Shim, J.Y., Kim, J.I., Choi, D.I., et al. (2022). Hippocampal engram networks for fear memory recruit new synapses and modify pre-existing synapses in vivo. Curr Biol.

Liu, X., Ramirez, S., Pang, P.T., Puryear, C.B., Govindarajan, A., Deisseroth, K., and Tonegawa, S. (2012). Optogenetic stimulation of a hippocampal engram activates fear memory recall. Nature 484, 381–385.

Matos, M., Bosson, A., Riebe, I., Reynell, C., Vallee, J., Laplante, I., Panatier, A., Robitaille, R., and Lacaille, J.C. (2018). Astrocytes detect and upregulate transmission at inhibitory synapses of somatostatin interneurons onto pyramidal cells. Nat Commun 9, 4254.

Megias, M., Emri, Z., Freund, T.F., and Gulyas, A.I. (2001). Total number and distribution of inhibitory and excitatory synapses on hippocampal CA1 pyramidal cells. Neuroscience 102, 527–540.

Muller, J.F., Mascagni, F., and McDonald, A.J. (2006). Pyramidal cells of the rat basolateral amygdala: synaptology and innervation by parvalbumin-immunoreactive interneurons. J Comp Neurol 494, 635–650.

Perea, G., Navarrete, M., and Araque, A. (2009). Tripartite synapses: astrocytes process and control synaptic information. Trends Neurosci 32, 421–431.

Ramirez, S., Liu, X., Lin, P.A., Suh, J., Pignatelli, M., Redondo, R.L., Ryan, T.J., and Tonegawa, S. (2013). Creating a false memory in the hippocampus. Science 341, 387–391.

Reilly, J.E., Hanson, H.H., and Phillips, G.R. (2011). Persistence of excitatory shaft synapses adjacent to newly emerged dendritic protrusions. Mol Cell Neurosci 48, 129–136.

Risher, W.C., Patel, S., Kim, I.H., Uezu, A., Bhagat, S., Wilton, D.K., Pilaz, L.J., Singh Alvarado, J., Calhan, O.Y., Silver, D.L., et al. (2014). Astrocytes refine cortical connectivity at dendritic spines. Elife 3.

Roy, D.S., Park, Y.G., Kim, M.E., Zhang, Y., Ogawa, S.K., DiNapoli, N., Gu, X., Cho, J.H., Choi, H., Kamentsky, L., et al. (2022). Brain-wide mapping reveals that engrams for a single memory are distributed across multiple brain regions. Nat Commun 13, 1799.

Schindelin, J., Arganda-Carreras, I., Frise, E., Kaynig, V., Longair, M., Pietzsch, T., Preibisch, S., Rueden, C., Saalfeld, S., Schmid, B., et al. (2012). Fiji: an open-source platform for biological-image analysis. Nat Methods 9, 676–682.

Sharma, V. (2018). ImageJ plugin HyperStackReg V5.6.

Stobart, J.L., Ferrari, K.D., Barrett, M.J.P., Gluck, C., Stobart, M.J., Zuend, M., and Weber, B. (2018). Cortical Circuit Activity Evokes Rapid Astrocyte Calcium Signals on a Similar Timescale to Neurons. Neuron 98, 726–735 e724.

Wable, G.S., Barbarich-Marsteller, N.C., Chowdhury, T.G., Sabaliauskas, N.A., Farb, C.R., and Aoki, C. (2014). Excitatory synapses on dendritic shafts of the caudal basal amygdala exhibit elevated levels of GABAA receptor alpha4 subunits following the induction of activity-based anorexia. Synapse 68, 1–15.

Yang, J., Wang, L., Yang, F., Luo, H., Xu, L., Lu, J., Zeng, S., and Zhang, Z. (2013). mBeRFP, an improved large stokes shift red fluorescent protein. PLoS One 8, e64849.

