## Supplemental Figures for "Astrocytic connection to engram neurons Increased after learning"

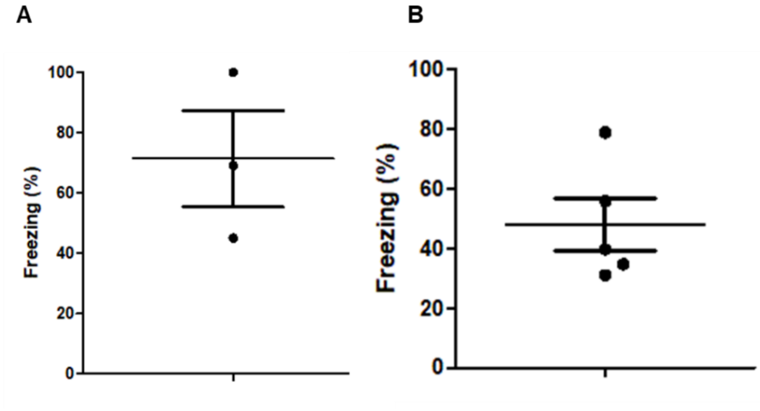


**Supplemental Figure 1. Freezing level for mice used for astrocyte-eGRASP analysis in Figure 4.**

(A) Freezing levels in mice used for astrocyte-eGRASP density analysis. n = 3. Data are represented as the mean ± SEM.

(B) Freezing levels in the mice used for dendritic spine morphology analysis. n = 5. Data are represented as the mean ± SEM.

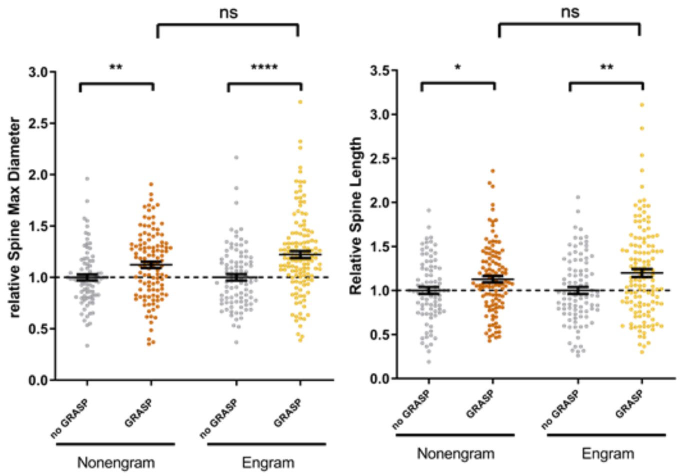


**Supplemental Figure 2. Increased spine head diameter and length are larger in dendritic spines with astrocyte-eGRASP.**

Dendritic spines with astrocyte-eGRASP show larger spine head diameter and spine length. Non-engram, no eGRASP spine, n = 84; non-engram, eGRASP spine, n = 114; engram, no eGRASP spine, n = 93; engram, eGRASP spine, n =131. Mann-Whitney two-tailed test. n.s., not significant, *$p < 0.05$, **$p < 0.01$, ****$p < 0.0001$. Data are represented as mean ± SEM.
